# Tau drives cell specific functional isolation of the hippocampal formation

**DOI:** 10.1101/2025.08.10.669580

**Authors:** Theodore J Zwang, Jasen Zhang, Rudy Gelb-Bicknell, Nina Wolf, Viktor Chanchykov, Haoyang Zhu, Ergina Agastra, Ava Devine, Andrew J. Holbrook, Rachel E. Bennett, Bradley T. Hyman

## Abstract

A major challenge in understanding Alzheimer’s disease is linking changes that occur across different biological scales. For example, how do changes in individual neurons build into widespread network disruptions? To address this, we used flexible mesh electronics to record neuronal activity for six months in ThyTau22 mice, a model of tauopathy that accumulates mutant human tau with age. Electrophysiology was recorded simultaneously from the hippocampus and entorhinal cortex of awake, behaving mice. At all ages we observed neuron-level, tau-driven silencing including ages without detectable tangles or cell-death. We found an unexpected phenomenon: neurons silenced by tau spontaneously recover individual firing patterns, yet these neurons fail to regain normal network interactions. Thus, as the animals age, disrupted network-level activity emerges. Specifically, we observe a global decrease in excitatory interactions and a breakdown in gamma-band coherence, which is particularly disrupted between the entorhinal cortex and hippocampus. These observations reveal a temporal relationship between neuronal silencing and impaired network connectivity, which also contributes to a progressive disruption in the excitatory/inhibitory balance. This ultimately disconnects viable entorhinal-hippocampal connections, physiologically isolating the hippocampus. Importantly, this network dysfunction is not driven by neuron loss, but by the failure of neurons to re-establish proper network interactions after silencing. This reveals a previously unrecognized mechanism by which mutant tau can destabilize neural systems. Further, these experiments indicate that a therapeutic window may exist where neuronal function and network activity might still be restored prior to irreversible degeneration.

## Introduction

The characteristic memory impairment observed in Alzheimer’s disease (AD) is strongly associated with tau related lesions, which are widely believed to cause the dysfunction of individual neurons and are correlated with neuronal loss across brain regions. Among the earliest and most vulnerable regions are the entorhinal cortex and hippocampus, which are essential for memory formation and retrieval. Critically, the projection neurons of the entorhinal cortex that give rise to the perforant pathway to the hippocampus accumulate tau pathology at the earliest disease stages.

Corresponding lesions in the CA1 and subiculum, an area that provides hippocampal-entorhinal and hippocampal-cortical output, is subsequently observed. These neurons are lost as the disease progresses, which is an irreversible event that effectively isolates the hippocampus from the cortex and correlates with memory impairments in Alzheimer’s disease^1^. Yet whether it is the tangles, the associated neuronal loss, or the presence of recently implicated misfolded oligomeric species of tau that leads to neural systems breakdown in Alzheimer’s disease is unknown.

Key to this problem is the difficulty of observing the integrity of both individual neurons and interconnected neural networks over time scales relevant to neurodegenerative disease. Tau can affect the firing activity of individual neurons and their synaptic connections with other neurons, which makes this problem ideal to study with longitudinal electrophysiology^2–4^ To better understand how the functional impacts of tau protein evolve over time within this memory circuit, we chose to employ a new generation of flexible mesh electronics. This technology has several key advantages, overcoming limitations of prior work. For one, it is minimally disruptive to the brain, mimicking the size and shape of neurons, which reduces damage to blood vessels during implantation and does not induce a glial response^5^. In turn, this allows stable recording from hundreds of neurons at a time over periods longer than 6 months, permitting the study of chronic neurodegenerative disease states^6,7^. By implanting electrodes in both the entorhinal cortex and hippocampus we can evaluate disrupted activity on multiple scales ranging from individual neuron dynamics to correlated neuron pairs and broader regional coherence patterns.

By combining this technology with a mouse model that develops tau pathology with age, ThyTau22^8^, we were able to observe how the function of individual neurons changes over time and how dysfunction in a subset of neurons impacts network activity. Importantly, these network-level disruptions occur at early ages and progress over months, preceding the formation of neurofibrillary tangles and occurring independently of neuronal death at the ages examined. These results implicate early, widespread cellular neurophysiological changes as a consequence of tau pathology. Neural system disruption and isolation of key memory related structures thus appear to be an emergent property of these individual neuron-level changes, perhaps in addition to, or instead of, the loss of neural system integrity consequent to neuronal death.

## Methods

### Design and fabrication of electronics

The overall fabrication approach of mesh electronics is similar to previous reports, but the designs used in this work are distinct and tailored to study of the brain regions of interest^5^. The designs of these mesh electronics are summarized in Supplementary Table 1, with the changes focused on incorporating 32-channel probes which are distributed across different lengths to optimally sample neurons within the hippocampus or entorhinal cortex.

The key fabrication steps for mesh electronics are as follows: (1) A 100-nm-thick Ni sacrificial layer was thermally evaporated (Sharon Vacuum Co.) onto a 4-inch diameter Si wafer (P/Boron, 0.001-0.005 Ω cm, 285 nm thermal oxide, Silicon Valley Microelectronics). (2) Negative photoresist SU-8 2000.5 (Kayaku) was mixed with Lissamine rhodamine B ethylenediamine (RhBen, Thermo Fisher Scientific) at a concentration of 10 µg/ml and placed in the dark at room temperature for 3 days.

This mixture was then centrifuged at 10,000 rpm for 3 minutes and supernatant collected/solids discarded prior to use. The SU-8 was then spin coated on top of the Ni layer at 4000 rpm, pre-baked sequentially at 65°C for 1 min and 95°C for 4 minutes, then patterned by photolithography with a mask aligner (SUSS MA6 mask aligner, SUSS MicroTec) using an i-line (365 nm wavelength) dose of 100 mJ/cm^2^. After photolithography, the wafer was post-baked sequentially at 65°C for 3 min then 95°C for 3 minutes. (3) The SU-8 photoresist was then developed in SU-8 developer (Kayaku) for 1 minute, rinsed with isopropanol, then dried with N_2_ and hard baked. Hard baking involved putting the wafer on a room temperature hot plate, ramping the temperature 1°C/min to 185°C, holding at 185°C for 60 minutes, then turning off and letting hot plate cool to room temperature. (4) The wafer was then spin-coated with LOR3A resist (Kayaku) at 4000 rpm and baked at 180°C for 5 minutes.

The wafer was cooled to RT then Shipley 1805 positive photoresist (Microposit, Dow Chemical Company) was spin coated onto the wafer at 4000 rpm and baked at 115°C for 1 minute. The positive resist was patterned by photolithography with h-line (405 nm wavelength) dose of 50 mJ/cm^2^) then developed in MF-CD-26 (Microposit, Dow Chemical Company) for 60 seconds. (5) A 10-nm-thick Ti layer and 100-nm-thick Au layer were sequentially deposited by electron beam evaporation (Denton Vacuum), followed by a liftoff step in Remover PG (Kayaku) with gentle shaking to remove the unpatterned gold from the surface and leave only the Au interconnect lines on the wafer. (6) Steps 4 and 5 were repeated for photolithography patterning, deposition of 3nm Ti then 50nm Pt on the recording electrodes, and liftoff of unpatterned metal. (7) Steps 2 and 3 were repeated for photolithography pattern of the top SU-8 layer, with care taken to ensure that the Au interconnect lines were completely insulated by the SU-8. The hard baking step was done ramping up to a temperature of 195°C, which ensured the two SU-8 layers were properly fused. (8) The mesh electronics were then cleaned with oxygen plasma at 40 ppm O_2_, 50 W, for 30 seconds (SCE 106 Quartz Barrel Plasma System, Anatech) and transferred to a Ni etchant solution (40% FeCl_3_:38% HCl:H_2_O mixed at a ratio of 1:1:20) until mesh was released from the surface, approximately 1-3 hours. Released mesh were immediately transferred to deionized water to rinse, then stored in phosphate buffered saline.

### Vertebrate animals

Adult male and female ThyTau22 mice (MAPT G272V, MAPT P301S) and littermate wild-type controls were used for this study^8^. Mice were bred in-house by crossing C57BL6 and heterozygous ThyTau22 mice. Mice were housed in a 12-hour reverse light/dark cycle and were provided food and water ad libitum. At 3 months of age the mesh were implanted (n=3 mice per genotype). All mice used in experiments had two successful implants with >80% connected channels in the same hemisphere. Additional naïve mice were used for histology experiments to evaluate tau and neuron loss at 3, 4, 5, 6, 9, 12, and 24 months of age (n=6 per genotype per age, with equal numbers of male and female mice).

All procedures were approved by the Institutional Animal Care and Use Committee of Massachusetts General Hospital. The animal care and use program at Massachusetts General Hospital meets the requirement of federal law and NIH regulations and are accredited by the American Association for Accreditation of Laboratory Animal Care.

### Mesh implantation surgery

Implantation of mesh electronics into live mice was performed using a stereotaxic injection method as described previously and reproduced here for completeness. All tools (Fine Science Tools) were autoclaved and bead-sterilized before use. Mesh electronics were sanitized with 70% ethanol followed by rinsing in sterile deionized water and sterile PBS prior to injection, then loaded into sterile glass capillaries with an inner diameter of 300 μm and outer diameter of 400 μm (Produstrial). Mice were anaesthetized in a chamber of 5% isoflurane, then moved to a head holder with a nose cone that provided passive 1-2% isoflurane exposure. A homeothermic blanket was set to 37°C and placed under the anesthetized mouse and the mouse was placed in a stereotaxic frame equipped with two ear bars and nose cone. Eye lubricant was applied to protect the eyes throughout the operation. Hair was removed with an electric razer and Betadine surgical scrub (Purdue) was applied to sanitize the scalp. A sterile scalpel was then used to make an incision along the sagittal sinus and the skin was resected to expose the skull.

3D printed plastic or titanium platform was attached to the skull via metabond dental cement (Purkell) and small screws leaving the sites above the hippocampus and entorhinal cortex visible. Using dental cement alone without screws increased likelihood of mechanical failure and detachment of the platform after several weeks or months of experiments.

Holes in the skull were drilled above the hippocampus and entorhinal cortex of the same hemisphere, as well as the cerebellum, then covered with surgical gelfoam soaked in saline. Ground screw was placed in the hole above the cerebellum. Mesh electronics were then slowly inserted using the glass capillary tube to the following coordinates, using the controlled injection method reported previously^5^: Mesh 1 was implanted anteroposterior −2.30 mm, mediolateral 1.75 mm, and dorsoventral 2.5 mm. Mesh 2 was implanted anteroposterior −4.70 mm, mediolateral 2.55 mm and dorsoventral 3.50 mm. Electronics were extruded from the glass capillary tube with a slow flow of sterile saline via syringe pump (PHD 2000, Harvard Apparatus) at 5ml/h for total injection volume <20µl, while capillary tube was retracted. The connection portion of the mesh electronics were then attached to a flat flexible cable (FFC, Premo-Flex, Molex) as described previously. After attachment, the connection between the mesh electronics and flat flexible cable were sealed using UV-curing glue (Visabella) upon UV exposure for 4 seconds, and further sealed with dental cement.

After surgery, each mouse was placed in a cage on a 37°C heating pad until ambulatory. The activity of the mouse was monitored until it fully recovered from anaesthesia, which typically occurred in <30 minutes. Liquid Tylenol was provided in the water, and Buprenorphine was given subcutaneously at a dose of 0.05 mg/kg every 12 hours for 72 hours post-surgery. Animals were housed individually after surgical procedures to minimize damage to electronics. All experiments were performed following a minimum 14 days of recovery.

### Chronic electrophysiological recording from awake and restrained mice

All experiments were done during the room dark cycle. To motivate the rodents, we used a water restriction. To start water restriction, over a 7-day period the water provided to the mice was decreased from 75 ml/kg to a final volume of 40m l/kg by 5 ml/kg increments per day. If no behavior was performed in a given day, mice were provided 40 ml/Kg water in cage dispensed with a calibrated pipette into a plastic petri dish. If the mouse had behavioral training or testing, then the amount of water received during training was calculated from the number of rewards and the volume of water per reward (∼4 µl). This amount was subtracted from the 40 ml/Kg and the mouse received the rest of the water in its cage dispensed as previously described after the task. This daily allotment was given at least 1 hour after testing so animals did not associate this water with the end of their behavioral task. The mice were monitored daily for signs of dehydration and weighed to ensure weight >85% starting weight. Mice have ad libitum access to dry food at all times in their cage. This water restriction only began after mice fully recovered from surgery.

If excessive weight loss (<80% from baseline) was found, mice were returned to ad lib water. If weight was not recovered upon return to ad lib water, mice were removed from study. We set exclusion criteria that if abnormal weight loss reached <75% from baseline or age-matched controls during water restriction, they would be removed from study, but this did not occur during our experiment. No significant health concerns were observed as a result of this approach.

Electrophysiology was recorded from mice while they were placed in a virtual reality system based on the design from the Harvey lab^9^. Mice were placed on top of a ball with two metal posts on the right and left of the mouse head used for restraint. The head platform installed during surgery has a hole on each side that can be fastened to these posts. This fixes the visual field of the mouse facing the virtual reality screen, but allows for full movement of the rest of the body and limbs on the ball.

During the first week, mice were trained for exploratory licking with 3x 10-minute sessions on different days in a training maze. Mice were given a 4 µl water reward whenever they licked the sensor, regardless of position. For each testing week (**Fig 1a**), the mice would be placed in a 110 cm long linear maze with a 20cm long lick-reward zone. When the mouse reached the end of the maze it would be teleported to the start. Mice were recorded for 15 minutes each test week, with the first 10 minutes having the VR system on and the last 5 minutes having the screen turned off.

**Figure 1.**
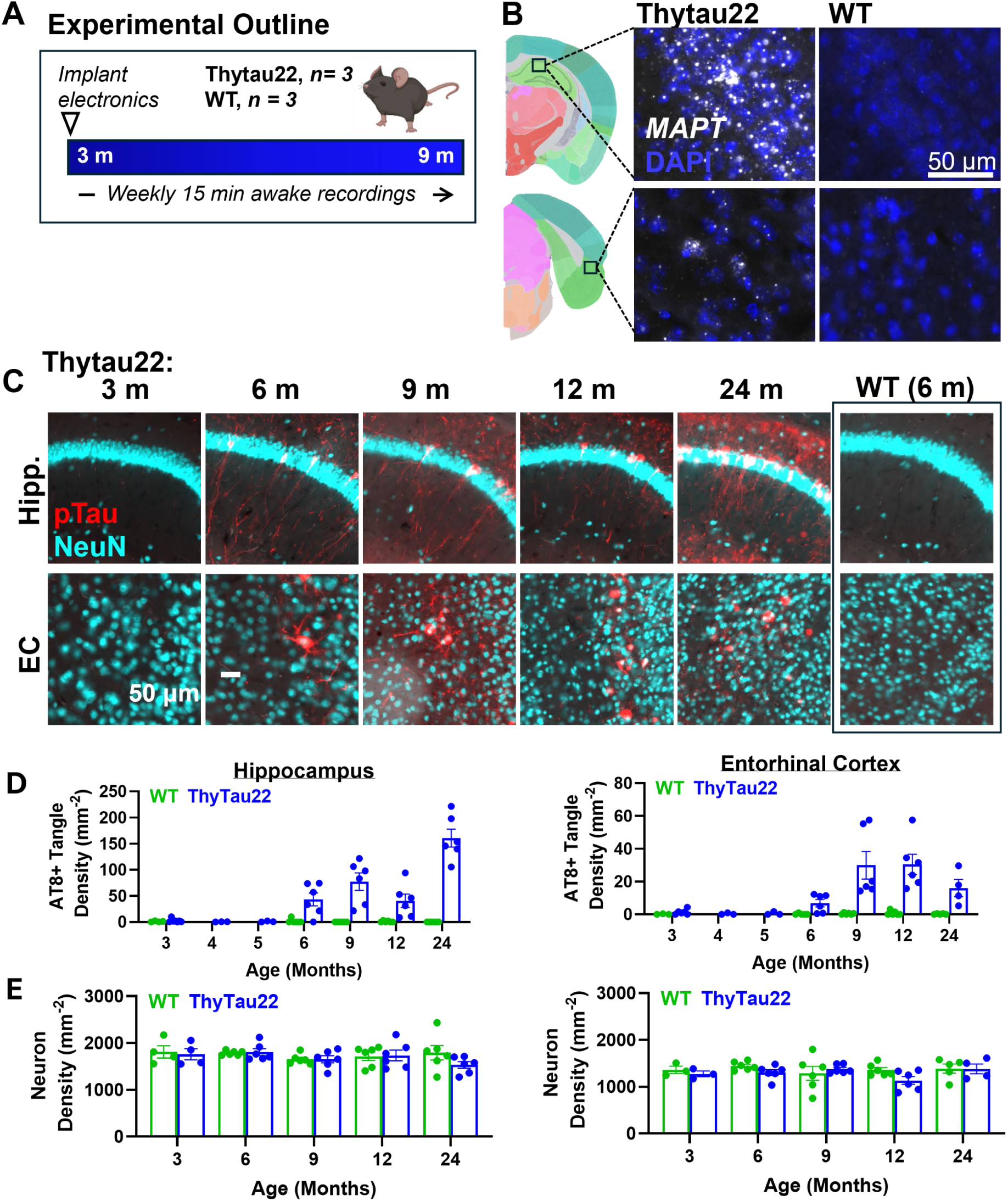
Thytau22 mice express human Tau and accumulate neurofibrillary tangles over time. (A) Experimental outline. Created with BioRender.com (B) *MAPT* in situ hybridization showing expression of human tau neurons throughout the entorhinal cortex and hippocampus of Thytau22 mice but not in the wild-type control. Diagram created using Allen Mouse Brain Atlas^33^. (C) Neurofibrillary tangles of phosphorylated tau (AT8 antibody, red) accumulate with age (m = months) in neurons (NeuN antibody, cyan) across the hippocampus and entorhinal cortex. (C) Quantification of AT8+ neurofibrillary tangle accumulation and (D) neuron density in the hippocampus and entorhinal cortex.

### Analysis of electrophysiology

Analysis of electrophysiology recordings was done using previously published workflow used successfully with flexible mesh electronics that involves loading the data into Matlab (2023b) and running voltage-time data in WaveClus3 to match individual units of neurons^5^. Spike times were extracted as a vector and used in the following analyses.

### Spike train preprocessing and cross-correlation analysis

Spike trains were generated at millisecond resolution from recorded spike times and stored in individual MATLAB files per neuron. These spike trains were concatenated across experimental sessions and grouped into non-overlapping bins, each encompassing four consecutive sessions (approximately four weeks). Spike train cross-correlograms were computed for each unique neuron pair:

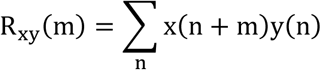

where x(n) and y(n) represent the binary spike train sequences of neuron pairs and m denotes the lag (in ms).

### Estimation of baseline coincidence rates using a Poisson model

Baseline coincidence rates were estimated from randomly shifted vectors using spike counts from lag intervals spanning ±10 to ±50 ms. Counts within these intervals were pooled to compute the mean baseline coincidence rate (λ), assuming spike coincidences follow a homogeneous Poisson distribution:

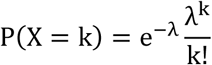

where k is the observed spike count, and λ is the mean count. Epochs were excluded from analysis if the expected baseline coincidences within the one-sided central window fell below two events, calculated as:

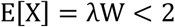

where W is the one side of the central window (5 ms total). Epochs were also excluded if there was a large excitatory peak precisely at 0ms, indicating a high probability of a duplicated unit.

### Detection of significant correlated pairs

Correlated pairs were probed within a central lag window of ±5 ms around zero lag. For neuron pairs recorded from the same electrode, ±2 ms around zero were excluded to mitigate spike collision artifacts. The total spike count (x) within the remaining central window was summed and compared against the expected sum under the Poisson model (λW). Statistical significance was assessed by converting Poisson-derived p-values to z-scores using the inverse normal transformation:

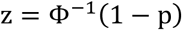

where Φ^−1^ denotes the inverse cumulative distribution function (CDF) of the standard normal distribution, and the p-value p was computed as:

- For excitation (observed counts > expected):

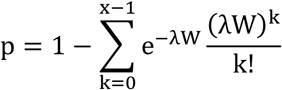

- For inhibition (observed counts < expected):

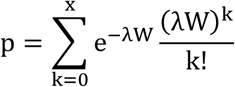

In cases where numerical underflow prevented exact calculation, Stirling’s approximation was applied to approximate the z-score:

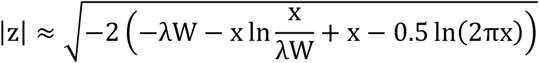

Results were separately recorded for positive and negative lags.

### Parallelization and I/O optimization

Analyses were parallelized using multiple MATLAB workers on a high-performance computing cluster. Neuron pairs were evenly distributed among workers via a custom function, and spike train data were dynamically loaded into memory using a least-recently-used caching strategy to optimize memory utilization and reduce disk I/O. All analyses were conducted in MATLAB R2024b.

### Statistical methods

Behavioral, spiketrain, and local field potential data were collected from three WT and three ThyTau22 mice in weekly experimental sessions. Data collection began when the mice were 14 weeks of age and continued until 38 weeks of age.

For each neuron, we calculated the firing rate as the number of spiketrain events divided by the length of the experimental session. Neurons that had zero spiketrain events for a given week were labeled as silent for that week. We then investigated the transience of silent weeks by calculating the average amount of consecutive silent weeks before a neuron reactivates, and this average was reported for each mouse across all its neurons. We performed Markov chain analysis, which captures the transience of a silent week by computing the empirical probability of a silent week being followed by a non-silent week.

To show silencing trends over time, we used logistic regression to model the proportion of silent neurons over time, and we also plotted a Kaplan Meier curve to visualize the time to a neuron’s first silent week.

Abnormal firing rates are defined as a firing rate that fell outside of the interquartile range. To capture the predictive nature of a neuron experiencing a silent week, we calculated the proportion of firing rates that were abnormal leading up to the first silent week of a neuron for each mouse.

Figures 1c, 1d, 2c, figures 4a-c were plotted and analyzed using Graphpad Prism (10.2), figure 4d was analyzed in Graphpad Prism (10.2) and plotted in excel. Figure hazard ratios for figure 4d were determined using chi-square analysis with the data in figure 4e-j. Figure 4e-j diagrams were created using SankeyMatic to represent accurate proportions. Figure 5a coherence was calculated using Brainstorm following their pre-processing and connectivity processes^10^. All other statistical analyses were conducted in R version 4.1.0, and all other figures were plotted with the ggplot2 R package.

**Figure 2.**
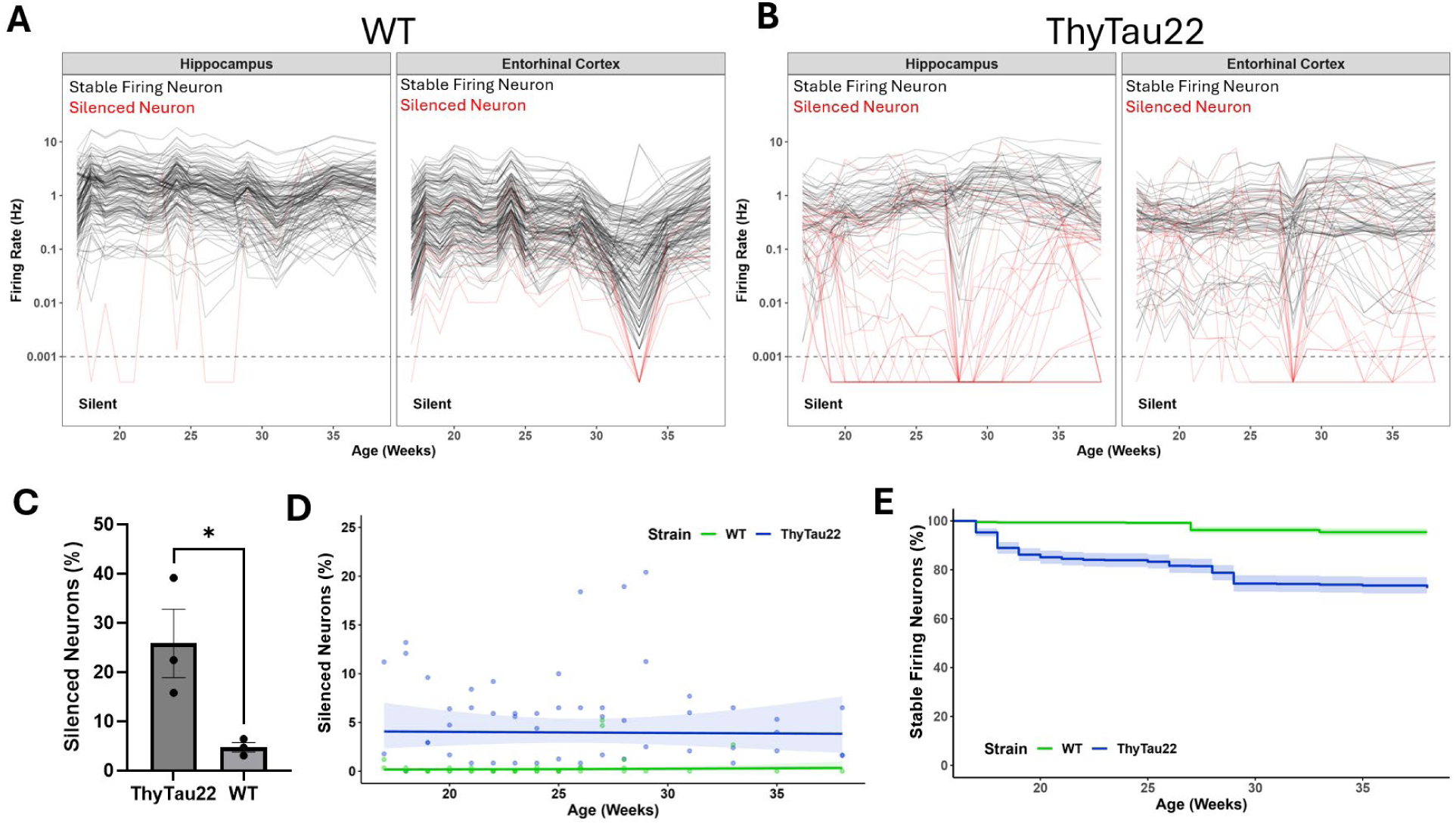
Most neurons show stable firing rates over months, but many neurons in ThyTau22 mice are temporarily silenced. **(A**.) Individual spike-sorted neuron firing rates are shown as each individual line for WT (**A**) and ThyTau22 (**B**) mice. Neurons with stable firing rates (black) and neurons that showed at least one silent week (red) are plotted over time. (**C**) Proportion of neurons that were defined as stable vs silenced in each mouse (n= 3 WT, 3 ThyTau22). (**D**) the number of neurons that are silent each week did not substantially change over time. (**E**) However, different neurons were silent across weeks leading to an increase in the number of neurons that have ever been silenced as ThyTau22 mice age.

### Tissue preparation and immunohistochemistry

At relevant ages, mice were perfused with phosphate buffered saline (PBS) and brains were extracted. The brains were fixed in 4% paraformaldehyde (PFA; Electron Microscopy Sciences) in PBS overnight then cryoprotected in 30% sucrose in PBS. Tissue was then sliced at 40 microns on a freezing microtome and stored in 30% glycerol in PBS until use. For immunolabeling the tissue was rinsed in PBS twice. Then the tissue was blocked with 5% bovine serum albumin (BSA, Sigma-Aldrich) in PBS with 0.25% Triton-X (Sigma-Aldrich) for 30 minutes. The tissue was then incubated with AT8 (Invitrogen, AB_223647) and/or NeuN antibody (1:5000; Abcam EPR12763) in blocking buffer overnight at 4C. Finally, tissue was rinsed with PBS twice. For HS84 labeling, the tissue was incubated with 1:1000 HS84 (from 5mg/ml stock solution; Sigma-Aldrich) in PBS for 30 minutes. The tissue was then rinsed 3x in PBS and cover slipped with Immumount.

### In situ hybridization

To examine human *MAPT* gene expression in tissue sections, we used the RNAscope Multiplex Fluorescent Reagent Kit v2 Assay (Advanced Cell Diagnostics) according to the manufacturer’s instructions for fixed frozen tissues and a HybEZ II hybridization system (Advanced Cell Diagnostics) with the following modifications. Prior to mounting free-floating sections on slides, tissues were rinsed in PBS and pre-treated with hydrogen peroxide (Advanced Cell Diagnostics) was performed in microcentrifuge tubes. Tissues were then placed on Superfrost Plus microscope slides (Fisher Scientific) and were then baked for 30 minutes in an oven at 60°C. Antigen retrieval steps included boiling in target retrieval solution (Advanced Cell Diagnostics) for 15 minutes and a 30 minute incubation with protease III (Advanced Cell Diagnostics). The human *MAPT* probe was applied (Hs-MAPT, Advanced Cell Diagnostic #408991) and detected with TSA Plus Cy3 (Akoya Biosciences, #NEL744E001KT). After completing the RNAscope steps, tissues were coverslipped with Fluoromount G with DAPI (Southern Biotech) and imaged using an Olympus VS120 slide scanning microscope.

## Results

### ThyTau22 mice accumulate tau pathology in the hippocampus and entorhinal cortex over months without overt neuronal loss

ThyTau22 (MAPT G272V, MAPT P301S) are a model for tau aggregation, which is a pathological hallmark of Alzheimer’s disease^11^. ThyTau22 express a four-repeat human tau isoform driven by the Thy1 promoter throughout the cortex and hippocampus (**Figure 1b**), and have been previously reported to accumulate a variety of tau-related neuropathological changes with age including tau hyperphosphorylation, neurofibrillary tangles^8^.

Coronal tissue sections of ThyTau22 mice were collected at 3, 6, 9, 12, and 24 months and immunohistochemically stained for neurons (NeuN) and phosphorylated tau (AT8) (**Figure 1c**). AT8-positive tau tangles generally appear sparingly at 3-5 months of age. At 6 months of age, we observe few tangles in the hippocampus and entorhinal cortex. At 9 months of age we observe substantial accumulation in both regions, which increases with age up to 24 months (**Figure 1d**). Comparing wild-type and Thytau22 mice revealed no significant change in neuron density in either the hippocampus or entorhinal cortex at any age (**Figure 1e**). We have previously observed mild neuron loss in this model with longitudinal imaging, and this discrepancy be attributed to low power to detect the small effect size in this cross-sectional analysis. Based on these data, we chose to record from mice starting at the pre-pathology 3 months age up to 9 months in order to capture electrophysiological changes due to the progressive accumulation of tau, prior to substantial tangle formation or neuronal loss.

### Flexible mesh electronics show stable recording from the same individual neurons over months in both ThyTau22 and WT mice

Two flexible electrophysiology probes were implanted within a single hemisphere of each mouse to record from the hippocampus and the medial entorhinal cortex. These regions were chosen due to their vulnerability to tau pathological deposition in Alzheimer’s disease and because the circuit is well conserved between species allowing us to meaningfully evaluate neuronal connectivity changes^12,13^. Recordings were made from 32 electrodes from each probe once per week for 15 minutes over 6 months while mice were awake and running on a ball that moved them through a linear track in virtual reality. Location of electronics was confirmed with histology (**Supplemental Figure 1**)

Clustering of waveforms revealed 778 individual neurons from WT mice and 659 neurons from ThyTau22 mice (**Figure 2**). 100% of neurons (778/778) from WT and 99% of neurons (653/659) from ThyTau22 mice were recorded throughout the 6-month window. Over this period there were no significant changes in the firing rates of neurons in the hippocampus and entorhinal cortex of WT mice (**Figure 2a**). In ThyTau22 mice, only 73% of neurons showed stable firing rates with the remainder showing abnormally low firing rates or were completely silent during at least one recording session (<0.001 Hz) (**Figure 2b,c**).

### Neurons undergo transient epochs of silencing throughout the lifespan of ThyTau22 mice

To further explore this neuronal silencing phenotype observed in tau mice, we next examined how single-neuron firing activity progressed with aging and tau accumulation. Across all weeks, an average of 4% of neurons are silent in ThyTau22 mice, compared to <1% in WT mice (**Figure 2d**). The percent of silent neurons per week did not change over time. We found 26% of all neurons are silent for at least one week and 97% of this subset of neurons show transient silencing, defined as neurons that fire <0.001 Hz during at least a single week but exceed this threshold in the following weeks (**Figure 2e**). Transiently silenced neurons were inactive for an average of 3.1 weeks before reactivating. Markov chain analysis shows that silenced neurons have a ∼39.04% (95% CI 35.22%-42.99%) probability of reactivating the following week, which was consistent in both the hippocampus (95% CI 33.66%-43.22%) and entorhinal cortex (95% CI 33.95%-47.17%).

Even before an epoch of silencing, individual neurons demonstrate an abnormal firing rate for several weeks (**Figure 3a**). Half of the neurons that become silent showed a significant decrease in firing rate 12 weeks prior to silencing in both the hippocampus (**Figure 3b**) and entorhinal cortex (**Figure 3c**). Neurons showed the ability to recover spontaneous activity, which was defined as an increase in firing rate until no statistically significant difference from the average firing rate of stable neurons. Silent neurons showed a slow recovery towards normal firing over the course of several weeks, though there were many neurons that never fully recovered their firing rates. These data indicate that tau can have a months-long trajectory of silencing and reactivation of neuronal firing. These trajectories were similar in both the hippocampus and entorhinal cortex.

**Figure 3.**
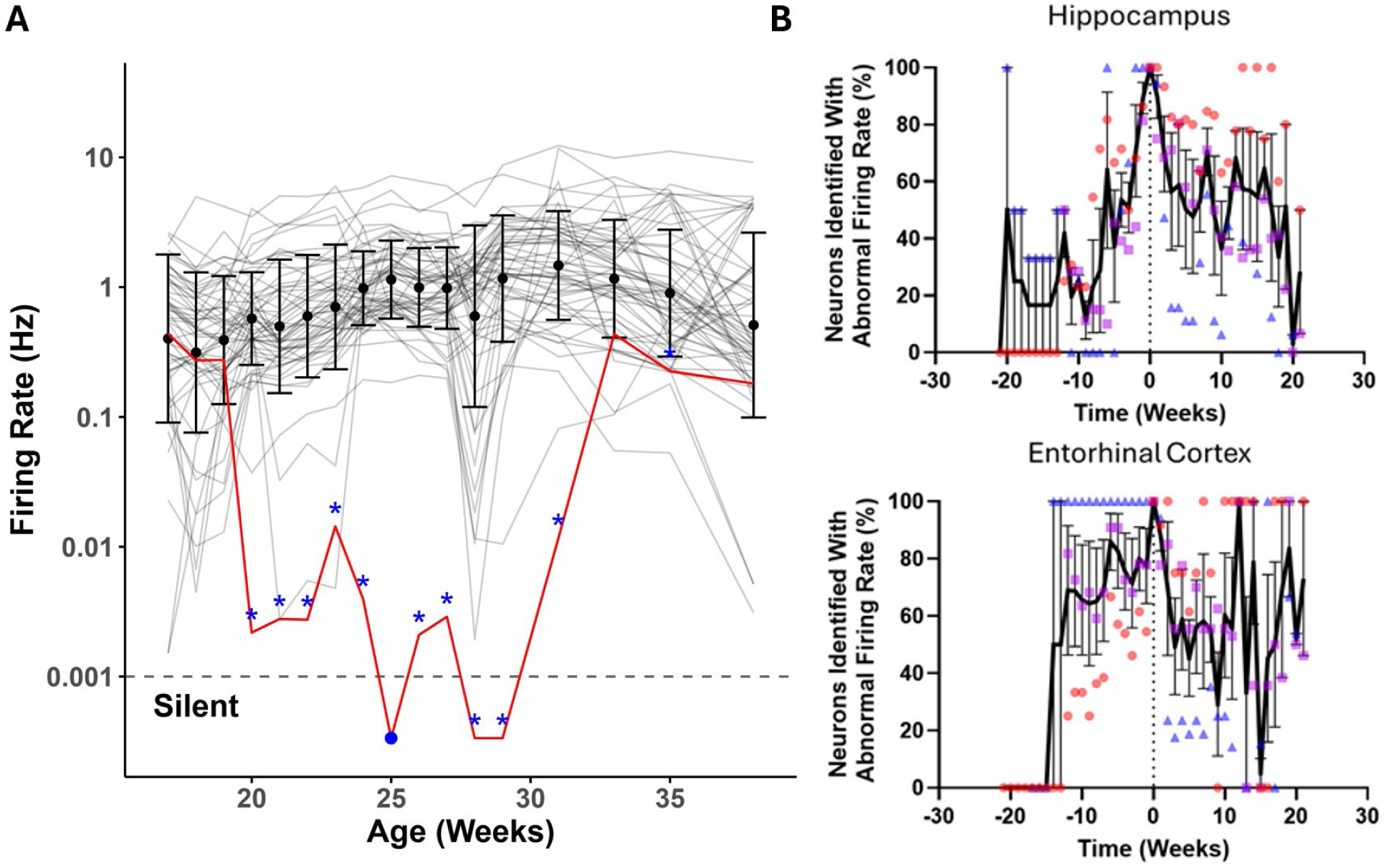
Plotting the trajectory of transient neuron silencing and recovery. (**A**) Neurons were defined as having abnormal firing (blue star) in any week that showed statistically significant firing rate decreases compared to the mean population of neurons. A neuron’s first silent week (blue circle) was defined as the first week with <0.001 Hz firing rate. We then plotted the percentage of neurons with abnormal firing rates relative to the time before or after their first silent week in both the (**B**) hippocampus and (**C**) entorhinal cortex. Each dot color represents a different mouse.

### Previously silenced neurons, even after recovering spontaneous activity, fail to reintegrate into circuits

ThyTau22 mice show substantial changes in the correlated activity of neurons over time, in stark contrast to the stability observed in WT mice. There is no significant difference in the percentage of neuron pairs that correlate within WT or within ThyTau22 mice, however the stability of correlated pairs is substantially decreased in ThyTau22 mice. In WT mice, 35.9% +/- 4.1% of excitatory and 43.5 +/- 8.9% of inhibitory correlated pairs are stable for at least 12 weeks but, in ThyTau22 mice, this decreases to 23.6% +/- 3.1% (two-tailed t-test, p = 0.038 vs WT) for excitatory pairs (**Figure 4a**) and 7.2+/- 1.4% (two-tailed t-test, p = 0.008 vs WT) for inhibitory pairs (**Figure 4b**). To isolate the effect of neuron silencing, we determined which correlations were between stable firing neurons and which correlations contained at least one silenced neuron, which was 18.2% +/- 9.6% of excitatory and 22.3% +/- 6.9% of inhibitory pairs (**Figure 4c**). When comparing only stable firing neurons in ThyTau22 mice to WT mice there was no statistically significant difference for the stability of excitatory (p = 0.86) or inhibitory (p = 0.25) correlated pairs (**Supplemental Figure 2**).

**Figure 4.**
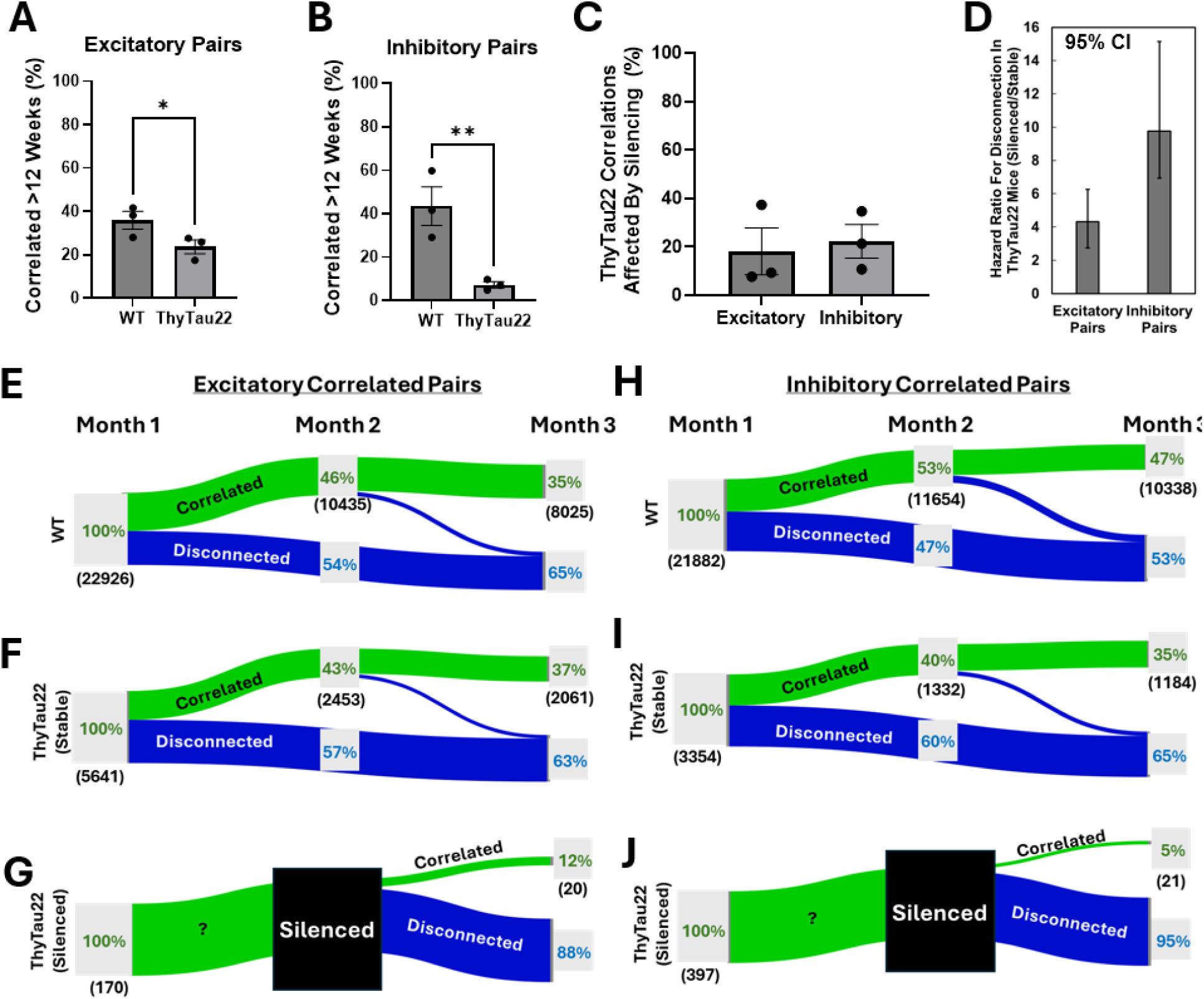
Silencing disrupts the correlated activity of neurons, even after neurons recover spontaneous activity. Spike train cross-correlograms were computed for each unique neuron pair to determine whether they represent an excitatory, inhibitory, or uncorrelated pair. Percent of (**A**) excitatory neuron pairs and (**B**) inhibitory neuron pairs which showed a statistically significant correlation for at least 12 weeks compared to all observed correlations for each mouse (n = 3 WT, 3 ThyTau22). Plot shows mean ± SEM. (**C**) Percent of all correlations in ThyTau22 mice that are affected by a neuron that is observed to be silenced. Plot shows mean ± SEM. (**D**) Hazard ratio for the disconnection of silenced vs stable neurons in ThyTau22 mice over three months. Plot shows mean and error bars are 95% CI. (**E-J**) Sankey diagram showing the change in connection for (**E**) excitatory and (**H**) inhibitory correlated pairs over three months in WT mice. The percentages in green represent the percent of statistically significant correlations, the black number in parenthesis are the number of correlations, and the blue percent represents the percent of disconnected correlations from pairs that were significantly correlated in the first month. ThyTau22 (**F**) excitatory and (**I**) inhibitory correlations between neurons that were stable over time, excluding any interactions with neurons that become silenced. ThyTau22 (**G**) excitatory and (**J**) inhibitory correlations between neurons that were correlated in the first month, silenced in the second month, then recovered activity in the third month to isolate the effect of silencing and reactivation on correlated pair stability. Only 6.3% of excitatory and 17.5% of inhibitory pairs that were affected by silencing met these criteria.

To isolate the effect of silencing on the stability of correlated pairs, we compared neurons that were correlated for one month, silent the following month, then recovered activity in the third month to the stability of correlated pairs from stable firing neurons in ThyTau22 mice and WT mice that were present during the first month of recording (**Figure 4e-j**). Only 6.3% (170) of excitatory and 17.5% (397) of inhibitory pairs that were affected by silencing at any time met these criteria. We observed a substantial reduction in the number neurons that were correlated with a previously silent neuron, even when the activity of the neuron recovered to baseline. We found excitatory neurons that recover from silencing were 4.3 times more likely to disconnect (Chi-square test, 95% CI 2.7-6.9, p < 0.001) and inhibitory neurons were 9.8 times (Chi-square test, 95% CI 6.3-15.2, p < 0.001) more likely to disconnect than neurons that had stable activity (**Figure 4d)**. Over time, we see an increase in the proportion of uncorrelated neurons in both the hippocampus and entorhinal cortex for ThyTau22 mice but not WT mice (**Supplemental Figure 3**). Altogether, these experiments show a decrease in correlated activity between neurons as ThyTau22 mice age that can be largely attributed to disconnections caused by the transient silencing of neurons.

### Local field potentials show abnormally high coherence followed by loss of coherence over weeks

We calculated local field potential coherence to quantify the frequency and amplitude of neuronal patterns of oscillating brain activity between pairs of electrodes at different frequency bands^14^. Local field potentials from microelectrodes have signals that originate within 250 µm of each recording electrode^15^, so we expect our electrode spacing of 60 µm should allow comparisons between different populations of neurons, especially for electrodes implanted in different brain regions. At early ages, Thytau22 mice show more coherent activity compared to WT in multiple frequencies for pairs of electrodes within the entorhinal cortex and pairs of electrodes hippocampus and entorhinal cortex (**Figure 5a**). WT shows stable coherence over the recording sessions at all frequencies. In stark contrast, coherence starts higher and decreases over time in ThyTau22 mice (**Figure 5b**). Gamma band activity shows the largest proportional change between ThyTau22 and WT. Coherence changes between electrodes across the hippocampus-entorhinal cortex, between electrodes within the hippocampus, and between electrodes within the entorhinal cortex all increased coherence at young ages followed by decreases over time in ThyTau22 compared to WT, but with substantial differences in magnitude and timing. The average hippocampal-entorhinal coherence for ThyTau22 starts substantially increased (101% +/- 69%), decreases to have similar coherence as WT by 16-18 weeks of age, then becomes less coherent than WT at 24 weeks of age and decreases to −48+/- 18% by 38 weeks of age. The average coherence between electrodes within the hippocampus show a substantial decrease over time, reaching values below WT from 24 weeks of age (−20.8% +/- 7.6%) onward. Electrodes within the entorhinal cortex start with increased average coherence (42% +/-15%) and show a slow decrease in coherence over the entire period, becoming statistically no different from WT by 23 weeks of age. Similar changes are reflected in other frequency bands in ThyTau22, but appear to decrease in magnitude as Gamma>Beta>Alpha>Theta>Delta (**Supplemental Figure 4**). The timing of inter-regional decoherence precedes tangle formation, but coincides with ages that show substantial silent neurons. Intra-regional decoherence follows weeks after and appears at similar age as tangle formation (∼24 weeks old). Altogether, these data suggest that progressive silencing of neurons with age may be involved in disconnecting neurons from network activity to ameliorate the abnormally high coherence in network activity.

**Figure 5.**
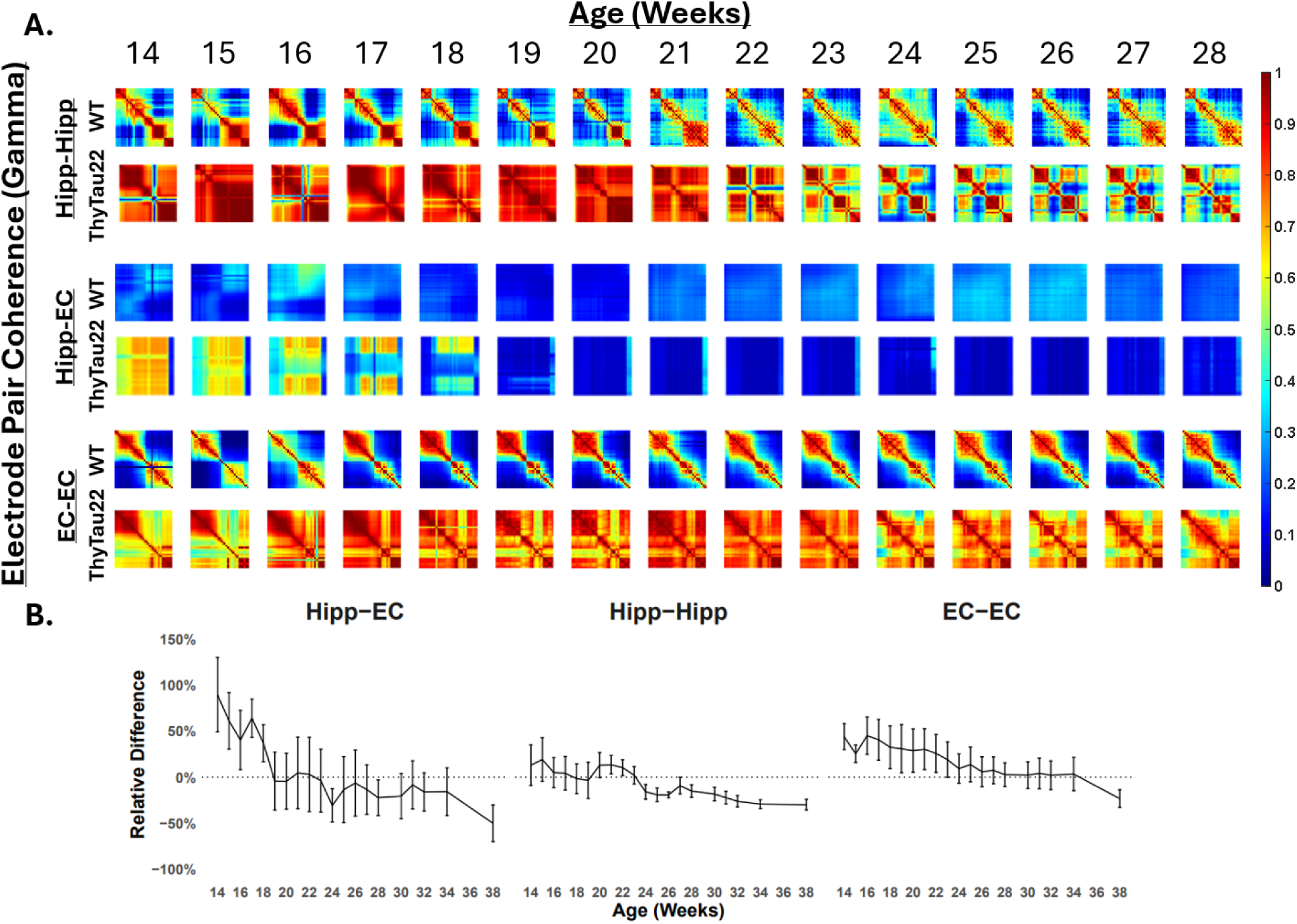
Electrode pair coherence over time. **A.** Change in coherence in gamma frequency (30-60Hz) for each electrode pair in the hippocampus (Hipp-Hipp), in the entorhinal cortex (EC-EC), and across both regions (Hipp-EC) for WT and ThyTau22 mice. Red represents more coherent and blue represents less coherent activity. **B.** Percent difference for the ratio ThyTau22 and WT gamma coherence (n=3 WT, 3 ThyTau22), shown as mean +/- SE.

## Discussion

While the role of tau inclusions and neuronal loss have been implicated in neural system dysfunction for decades, the current experiments surprisingly reveal a role for tau, without tangles or neuronal loss, in a complex and dynamic trajectory of electrophysiological changes that ultimately affects multiple levels of brain organization—from individual neurons to coordinated neuronal pairs to network-level local field potential coherence. Although we and others have previously described reduced average neuronal activity in tau expressing animals^4,16,17^, our longitudinal assessments now demonstrate that tau-induced neuronal silencing occurs in weeks-long epochs. This silencing, despite being stable in an individual neuron for weeks to months, is reversible, and reactivation often occurs. However, silencing appears to impact the stability of established neural circuits in which the silenced neuron had previously participated: plasticity events presumably occur, leading to loss of re-establishment of coordinated pairs of neurons even after resumption of firing. Indeed, as the number of individual neurons that undergo silencing grows over time, the stability of the neural system is ultimately impacted. Thus, fundamental physiological changes in individual neurons accumulate and lead to an emergent property of disruption of coherent firing patterns in the entorhinal cortex and hippocampus, reflected by age dependent changes in local field potentials and inter-regional loss of coherent signaling. This physiological change, even without neuronal loss or substantial tangle formation, likely contributes to the impairments in normal memory function observed in this strain^8,18,19^, which depends on the integrity of entorhinal-hippocampal connections.

Our recordings in awake, behaving mice over 6 months represent a substantial advancement over prior studies of tau pathology, which could not distinguish the same neurons or circuits across long times. Our results confirm previous observations of progressive decreases in coordinated activity of neurons^20^, but tracking individual neurons was essential for observing the trajectory of neuron silencing and its impact on coordinated activity. We found that at any given time point, only a small fraction of neurons were silent. Each week, roughly half of the previously silent neurons reactivated while a new group became silent. In studies using Neuropixels probes, such transient activity changes may have been misattributed to electrode drift and thus excluded from analysis^21,22^. However, our use of flexible mesh electronics minimizes drift and has previously been used to track individual neurons on months-to-years timescales, and the characteristic gradual reduction in firing rate followed by a slow recovery further supports a biological rather than artifactual explanation^5,7,23^.

Neuron silencing appears to be an early stage and key contributor leading to tau-related neural network dysfunction. Although many neurons recover spiking activity, their ability to engage in coordinated firing with other neurons remains impaired. Recovered neurons exhibited reduced pairwise correlations, indicating persistent disconnection from broader network activity. Interestingly, ThyTau22 LFP coherence was initially elevated and declined over time to be comparable to WT. The increased coherence observed here appears to have begun at ages younger than our earliest recording and is likely initiated by the 5-6 fold higher levels of 4R tau expressed in this line, which begins at birth and may impact development although no behavioral deficits are observed at these early ages^8^. Similarly early elevations in activity have previously been observed by EEG in a separate tau mouse line^24^. Conversely, tau knock-out lines have been found to be protective against seizures^25^. These patterns suggest that tau levels functionally alter neuronal activity and neuron silencing may be a response to restore dysfunctional networks. In tauopathies such as frontotemporal dementia and Alzheimer’s disease we speculate that this process may be dysregulated and trigger a maladaptive pruning process that attempts to stabilize network activity by disconnecting aberrant nodes. These results raise the possibility that the fundamental cellular lesion that impacts neural system integrity is, at least at the start, reflecting reversible physiological changes. Ultimately, though plasticity mechanisms, this leads to excitatory inhibitory imbalance, changes in local field potentials, and neural system isolation. Importantly, this loss of neural system integrity occurs without requiring frank tangle formation or overt neuronal loss.

The finding that neurons can recover activity and survive despite prolonged silencing suggests a potentially wide therapeutic window in which interventions could restore function before irreversible degeneration occurs. Previous observations of tau-associated reduction in activity or silencing of individual neurons were made in cross-sectional or in vitro models (e.g., anesthetized mice, brain slices, or cultured neurons), which precluded observation of spontaneous reactivation^16,17,26^. In contrast, our data show that neurons can recover firing activity, possibly by reducing or clearing intracellular tau accumulation, as suggested by studies where tau suppression can improve memory function and leads to a decrease in the number of silent neurons^16,27^.

Neither neuron loss or tangle formation are required for initiating neuron silencing and subsequent disconnection of neuronal networks that result from mutant human tau, which indicates that interventions may need to preserve functional connections independently of protecting neurons from death or other consequences of protein aggregation. Tau has been found to affect synaptic function and plasticity with consequences for memory^28,29^, but with some timing dependence for intervention efficacy ^30^. We speculate that sustained suppression of this tau effect may ultimately allow successful plasticity of remaining neurons at any stage of the disease, potentially restoring function even after the disease is established. Indeed, the initial studies of aged rTg4510 mouse (expressing P301L tau transgene) showed a robust recovery of behavior and electrophysiology after suppression of the tau transgene, despite the presence of substantial neuronal loss and tangles^27^.

In sum, this work links single-neuron and systems-level observations to tau pathology by demonstrating that early loss of correlated and network activity is initiated by neuron silencing, which may be a response to excessively coordinated brain activity. Additionally, our findings show that these silent neurons are capable of reactivation over time. This has important implications for interpreting noninvasive brain signals in human patients. Changes in LFP power may reflect the transient silencing, inactivity, or loss of neurons. Changes in LFP coherence may more accurately reflect the disruption in synaptic communication and circuit integrity that progresses from a state of excessive synchrony. Thus, coherence may serve as a more sensitive measure of tau-induced synaptic dysfunction, because it reflects the timing relationships between regions or neurons.

Related measurements of hypersynchronization have been observed in human studies, and appears closely related to progression from mild cognitive impairment to Alzheimer’s disease^31,32^. Our results suggest LFP coherence measures may be an important readout for anti-tau therapeutic trials and may be more sensitive than static imaging biomarkers of fibrillar tau in understanding the physiological impacts of potential therapeutics.

## Acknowledgements

We are grateful to our colleagues for their support and discussion. We thank Harvard’s Center for Nanoscale Systems for use of their facilities for electronics fabrication, and Massachusetts General Hospital’s Center for Comparative Medicine for their support in animal care. We thank Christopher D. Harvey and Noah L. Pettit for their help in building virtual reality systems.

---

**Supplemental Figure 1.**
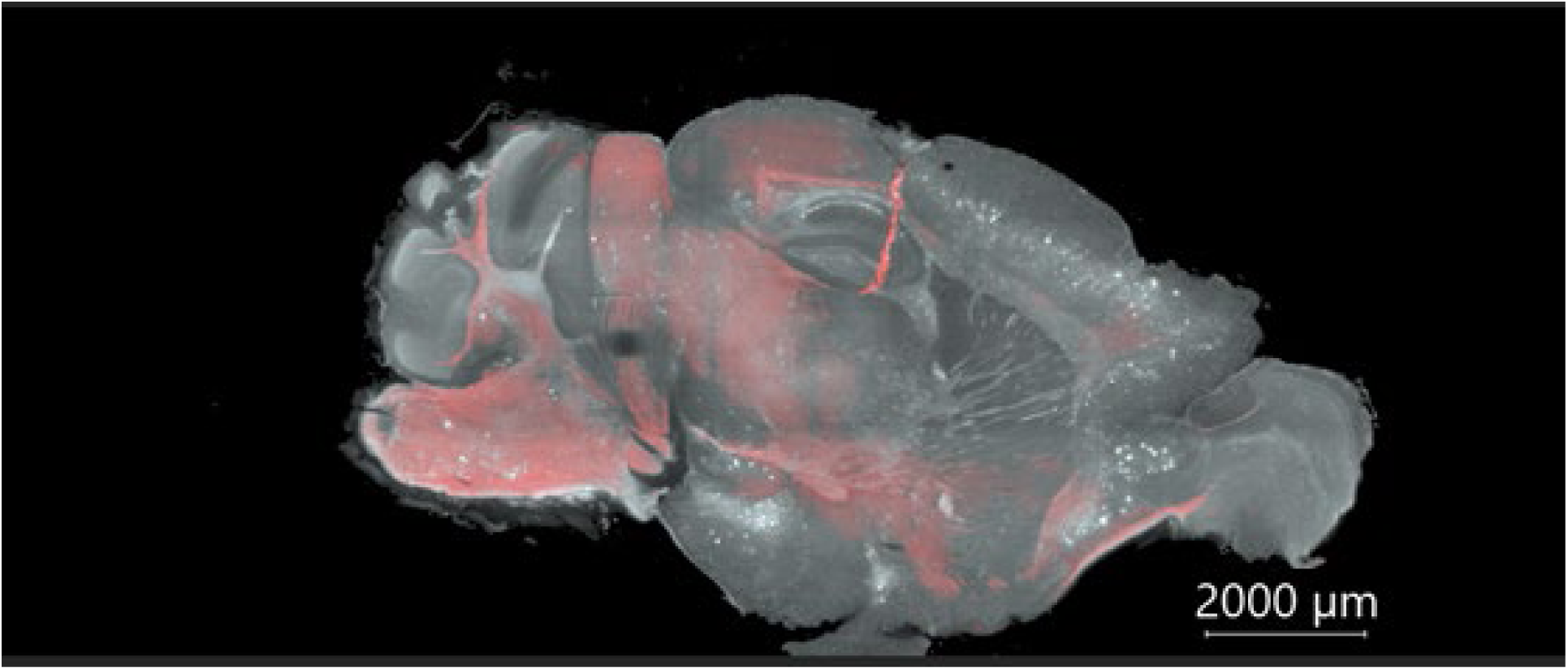
Location of electronics (red) implanted in the hippocampus of a ThyTau22 mouse stained with AT8 (white).

**Supplemental Figure 2.**
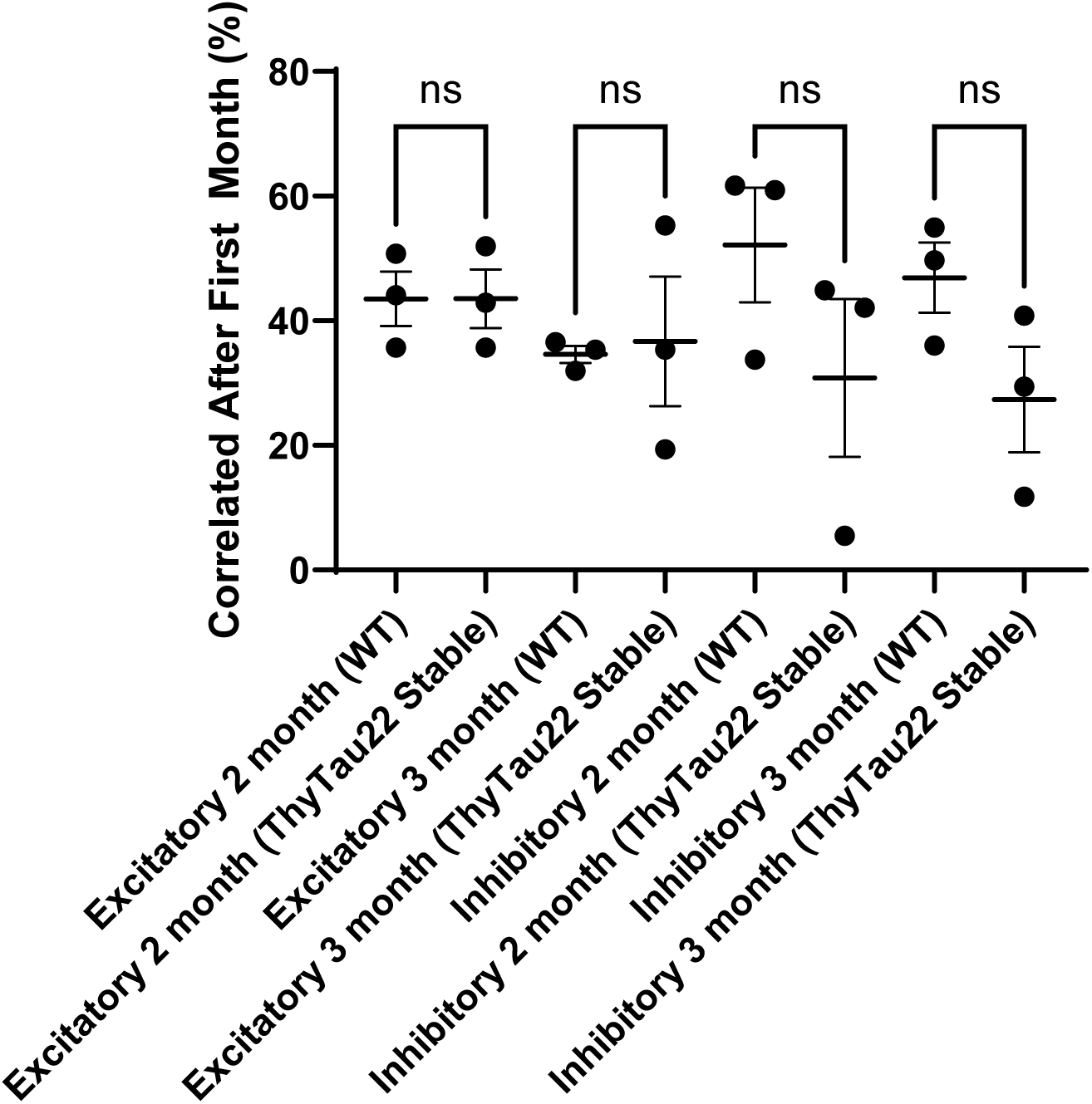
No difference in the stability of correlated neuron pairs after 2- or 3-months of recording when comparing WT and ThyTau22 stable-firing neurons.

**Supplemental Figure 3.**
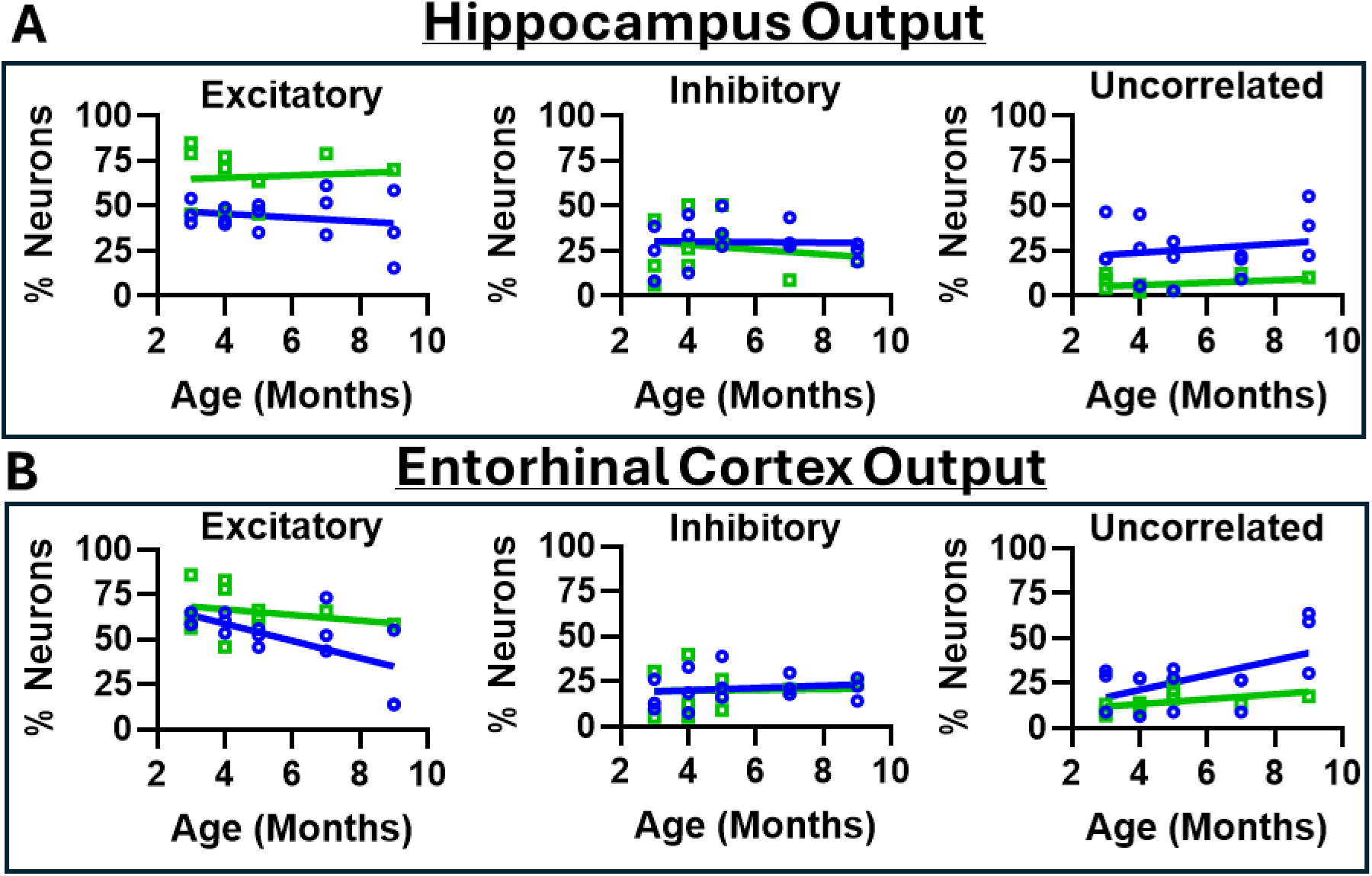
The number of neurons with excitatory output decreases over time in both the hippocampus and entorhinal cortex in Thy-Tau22 (blue) but not WT (green) mice. The number of neurons with inhibitory output do not change over time for either Thy-Tau22 or WT mice. Each shape represents the proportion of neurons from an individual mouse (n=3 WT, 3 ThyTau22). Values were normalized as a percentage of all neurons recorded in each individual mouse. These data show neurons with excitatory correlations are becoming uncorrelated rather than permanently inactivating or dying.

**Supplemental Figure 4.**
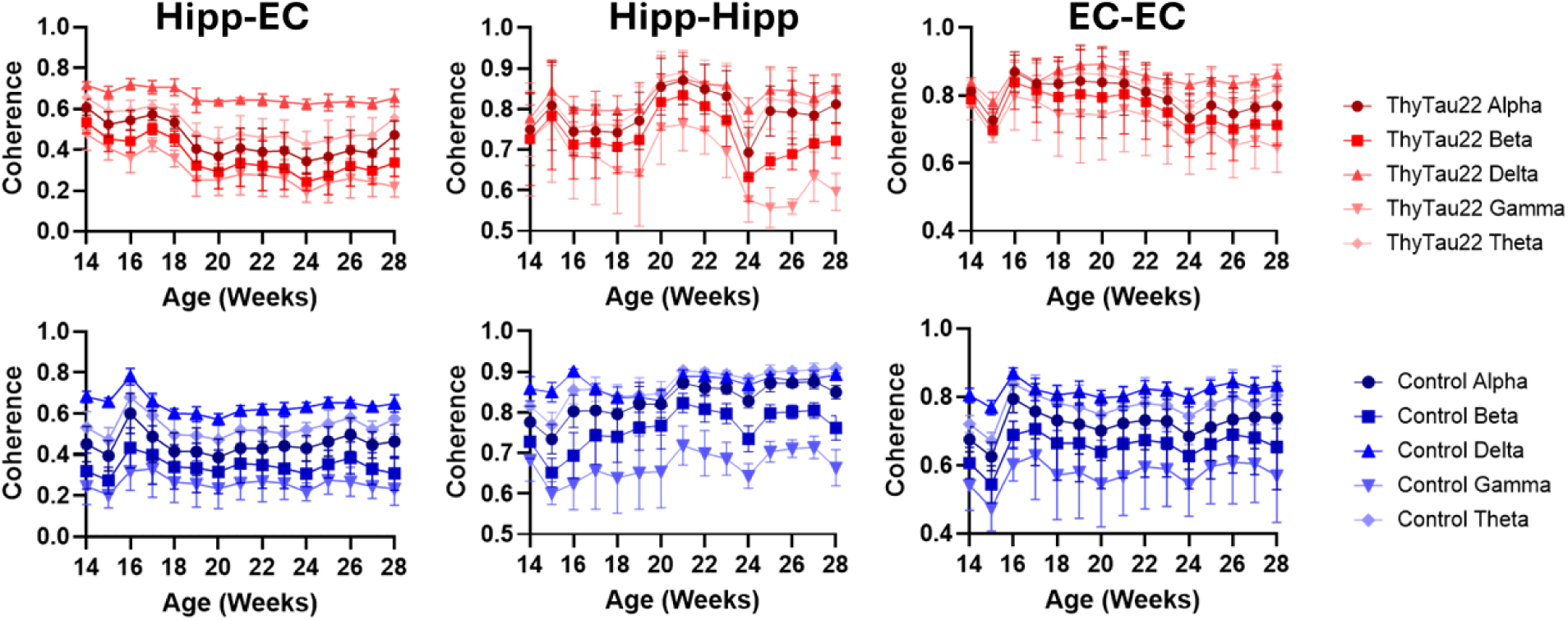
Average change in coherence across mice (n=3 WT, 3 ThyTau22) for different filtered frequency bands.

## References

1 Hyman, B. T., Van Hoesen, G. W., Damasio, A. R. & Barnes, C. L. Alzheimer’s Disease: Cell-Specific Pathology Isolates the Hippocampal Formation. Science 225, 1168–1170 (1984). doi:10.1126/science.6474172

2 Crimins, J. L., Pooler, A., Polydoro, M., Luebke, J. I. & Spires-Jones, T. L. The intersection of amyloid beta and tau in glutamatergic synaptic dysfunction and collapse in Alzheimer’s disease. Ageing Research Reviews 12, 757–763 (2013). 10.1016/j.arr.2013.03.002

3 Busche, M. A. Tau suppresses neuronal activity in vivo, even before tangles form. Brain 142, 843–846 (2019). 10.1093/brain/awz060

4 Menkes-Caspi, N. et al. Pathological Tau Disrupts Ongoing Network Activity. Neuron 85, 959–966 (2015). 10.1016/j.neuron.2015.01.025

5 Yang, X. et al. Bioinspired neuron-like electronics. Nature Materials 18, 510–517 (2019). 10.1038/s41563-019-0292-9

6 Fu, T.-M., Hong, G., Viveros, R. D., Zhou, T. & Lieber, C. M. Highly scalable multichannel mesh electronics for stable chronic brain electrophysiology. Proceedings of the National Academy of Sciences 114, E10046–E10055 (2017). doi:10.1073/pnas.1717695114

7 Fu, T.-M. et al. Stable long-term chronic brain mapping at the single-neuron level. Nature Methods 13, 875–882 (2016). 10.1038/nmeth.3969

8 Schindowski, K. et al. Alzheimer’s Disease-Like Tau Neuropathology Leads to Memory Deficits and Loss of Functional Synapses in a Novel Mutated Tau Transgenic Mouse without Any Motor Deficits. The American Journal of Pathology 169, 599–616 (2006). 10.2353/ajpath.2006.060002

9 Pettit, N. L., Yuan, X. C. & Harvey, C. D. Hippocampal place codes are gated by behavioral engagement. Nature Neuroscience 25, 561–566 (2022). 10.1038/s41593-022-01050-4

10 Tadel, F., Baillet, S., Mosher, J. C., Pantazis, D. & Leahy, R. M. Brainstorm: A User-Friendly Application for MEG/EEG Analysis. Computational Intelligence and Neuroscience 2011, 879716 (2011). 10.1155/2011/879716

11 Jack Jr., C. R., et al. Revised criteria for diagnosis and staging of Alzheimer’s disease: Alzheimer’s Association Workgroup. Alzheimer’s & Dementia 20, 5143–5169 (2024). 10.1002/alz.13859

12 Allen, T. A. & Fortin, N. J. The evolution of episodic memory. Proceedings of the National Academy of Sciences 110, 10379–10386 (2013). doi:10.1073/pnas.1301199110

13 Braak, H., Alafuzoff, I., Arzberger, T., Kretzschmar, H. & Del Tredici, K. Staging of Alzheimer disease-associated neurofibrillary pathology using paraffin sections and immunocytochemistry. Acta Neuropathologica 112, 389–404 (2006). 10.1007/s00401-006-0127-z

14 Bowyer, S. M. Coherence a measure of the brain networks: past and present. Neuropsychiatric Electrophysiology 2, 1 (2016). 10.1186/s40810-015-0015-7

15 Katzner, S. et al. Local Origin of Field Potentials in Visual Cortex. Neuron 61, 35–41 (2009). 10.1016/j.neuron.2008.11.016

16 Busche, M. A. et al. Tau impairs neural circuits, dominating amyloid-β effects, in Alzheimer models in vivo. Nature Neuroscience 22, 57–64 (2019). 10.1038/s41593-018-0289-8

17 Marinković, P. et al. In vivo imaging reveals reduced activity of neuronal circuits in a mouse tauopathy model. Brain 142, 1051–1062 (2019). 10.1093/brain/awz035

18 Van der Jeugd, A. et al. Progressive Age-Related Cognitive Decline in Tau Mice. Journal of Alzheimer’s Disease 37, 777–788 (2013). 10.3233/jad-130110

19 Lo, A. C. et al. Amyloid and Tau Neuropathology Differentially Affect Prefrontal Synaptic Plasticity and Cognitive Performance in Mouse Models of Alzheimer’s Disease. Journal of Alzheimer’s Disease 37, 109–125 (2013). 10.3233/jad-122296

20 McGregor, J. N. et al. Failure in a population: Tauopathy disrupts homeostatic set-points in emergent dynamics despite stability in the constituent neurons. Neuron 112, 3567–3584.e3565 (2024). 10.1016/j.neuron.2024.08.006

21 van Beest, E. H. et al. Tracking neurons across days with high-density probes. Nature Methods 22, 778–787 (2025). 10.1038/s41592-024-02440-1

22 Garcia, S. et al. A Modular Implementation to Handle and Benchmark Drift Correction for High-Density Extracellular Recordings. eneuro 11, ENEURO.0229-0223.2023 (2024). 10.1523/eneuro.0229-23.2023

23 Zhao, S. et al. Tracking neural activity from the same cells during the entire adult life of mice. Nature Neuroscience 26, 696–710 (2023). 10.1038/s41593-023-01267-x

24 Holth, J. K., Mahan, T. E., Robinson, G. O., Rocha, A. & Holtzman, D. M. Altered sleep and EEG power in the P301S Tau transgenic mouse model. Annals of Clinical and Translational Neurology 4, 180–190 (2017). 10.1002/acn3.390

25 DeVos, S. L. et al. Antisense Reduction of Tau in Adult Mice Protects against Seizures. The Journal of Neuroscience 33, 12887–12897 (2013). 10.1523/jneurosci.2107-13.2013

26 Opland, C. K. et al. Activity-dependent tau cleavage by caspase-3 promotes neuronal dysfunction and synaptotoxicity. iScience 26 (2023). 10.1016/j.isci.2023.106905

27 SantaCruz, K. et al. Tau Suppression in a Neurodegenerative Mouse Model Improves Memory Function. Science 309, 476–481 (2005). 10.1126/science.1113694

28 Ittner, L. M. et al. Dendritic Function of Tau Mediates Amyloid-β Toxicity in Alzheimer’s Disease Mouse Models. Cell 142, 387–397 (2010). 10.1016/j.cell.2010.06.036

29 Martinez, P. et al. Bassoon contributes to tau-seed propagation and neurotoxicity. Nature Neuroscience 25, 1597–1607 (2022). 10.1038/s41593-022-01191-6

30 Tang, S. J. et al. Fyn kinase inhibition reduces protein aggregation, increases synapse density and improves memory in transgenic and traumatic Tauopathy. Acta Neuropathologica Communications 8, 96 (2020). 10.1186/s40478-020-00976-9

31 Pusil, S. et al. Hypersynchronization in mild cognitive impairment: the ‘X’ model. Brain 142, 3936–3950 (2019). 10.1093/brain/awz320

32 Maestú, F., de Haan, W., Busche, M. A. & DeFelipe, J. Neuronal excitation/inhibition imbalance: core element of a translational perspective on Alzheimer pathophysiology. Ageing Research Reviews 69, 101372 (2021). 10.1016/j.arr.2021.101372

33. Allen Institute for Brain Science (2004). Allen Mouse Brain Atlas [dataset]. Available from mouse.brain-map.org. Allen Institute for Brain Science (2011).

